# Mental calculation speeds surpass known limits for high-level cognition

**DOI:** 10.1101/2025.05.28.656137

**Authors:** Bastian Wiederhold, Martin Stemmler, Andreas V.M. Herz

**Affiliations:** Bernstein Center for Computational Neuroscience and Faculty of Biology, Ludwig-Maximilians-Universität, Munich

## Abstract

While our senses transmit information at rates exceeding 10^6^ bit/s, high-level cognitive processing is thought to be much slower, on the order of 10 bit/s regardless of the task^1^. It is unclear, though, whether this limit holds when the human mind is challenged. To test how fast one can process abstract information, we analyzed mental calculations, using data from international competitions and record-setting attempts. We discovered scaling laws relating task complexity and completion speed. High information rates were sustained for minutes, peaking at above 215 bit/s for the shortest calculations, split into 125 bit/s for conscious perception and 90 bit/s for algorithmic execution. These rates are well in excess of previous estimates for high-level mental abilities.

---

In a recent study^1^, Zheng and Meister argued that the human mind possesses a universal information processing limit at around 10 bit/s, irrespective of the nature of the task. This rate might reflect the complexity of conscious decisions under ordinary circumstances, but prompts the question of what human beings are capable of doing when pushed to the limit.

For studying the bounds of performance, the field of mental calculation offers several advantages. There are standardized, established competitions and the visual input consists of plain decimal numbers, perfectly suited for reliable and quantitative studies. Contrary to other cognitive activities, the employed mental strategies are based on explicit algorithms, which allow us to precisely estimate the internal processing steps. Evidence suggests that expert mental calculation is a learned skill^2–6^ and, indeed, human “calculators” perform within the standard range in other cognitive tests^7,8^. Hence, mental calculation records inform us about human cognition as the result of practice and training, rather than the abilities of singularly gifted individuals.

We analyze the bit rates achieved in four broad competition categories: *addition, multiplication, calendar dates* and *square roots*. The combinatorial number of possible answers in these challenges precludes elaborate, memorized lookup tables, so competitors apply learned-by-rote arithmetic operations repeatedly in rapid succession. We account for mental calculation “tricks” and human working memory capacity, which dictates the competitors’ choice of algorithm, speeding them up or slowing them down, respectively. To capture the different cognitive dimensions involved, we compute the bit rates associated with three key steps in the process: reading, calculating, and answering. The sum of these rates falls off with task duration in a power-law fashion and peaks at more than 215 bit/s. This rate surpasses estimates for high-level cognition at least threefold^9–14^ or even more^1^.

## Calculation duration correlates with task entropy

Most mental calculations consist of three stages: transforming a visual input into a mental representation, executing the algorithm, and reporting the answer through motor activity as the output (Fig. 1 **A**). The complexity of a calculation is measured in bits (Shannon entropy), which is the logarithm of the number of possibilities, weighted by their likelihood, that need to be entertained at each stage. In competitions and record-setting attempts, error-free performances are often the only ones rewarded, so that the entropy rate bounds the calculators’ mental information rate from below.

**Figure 1.**
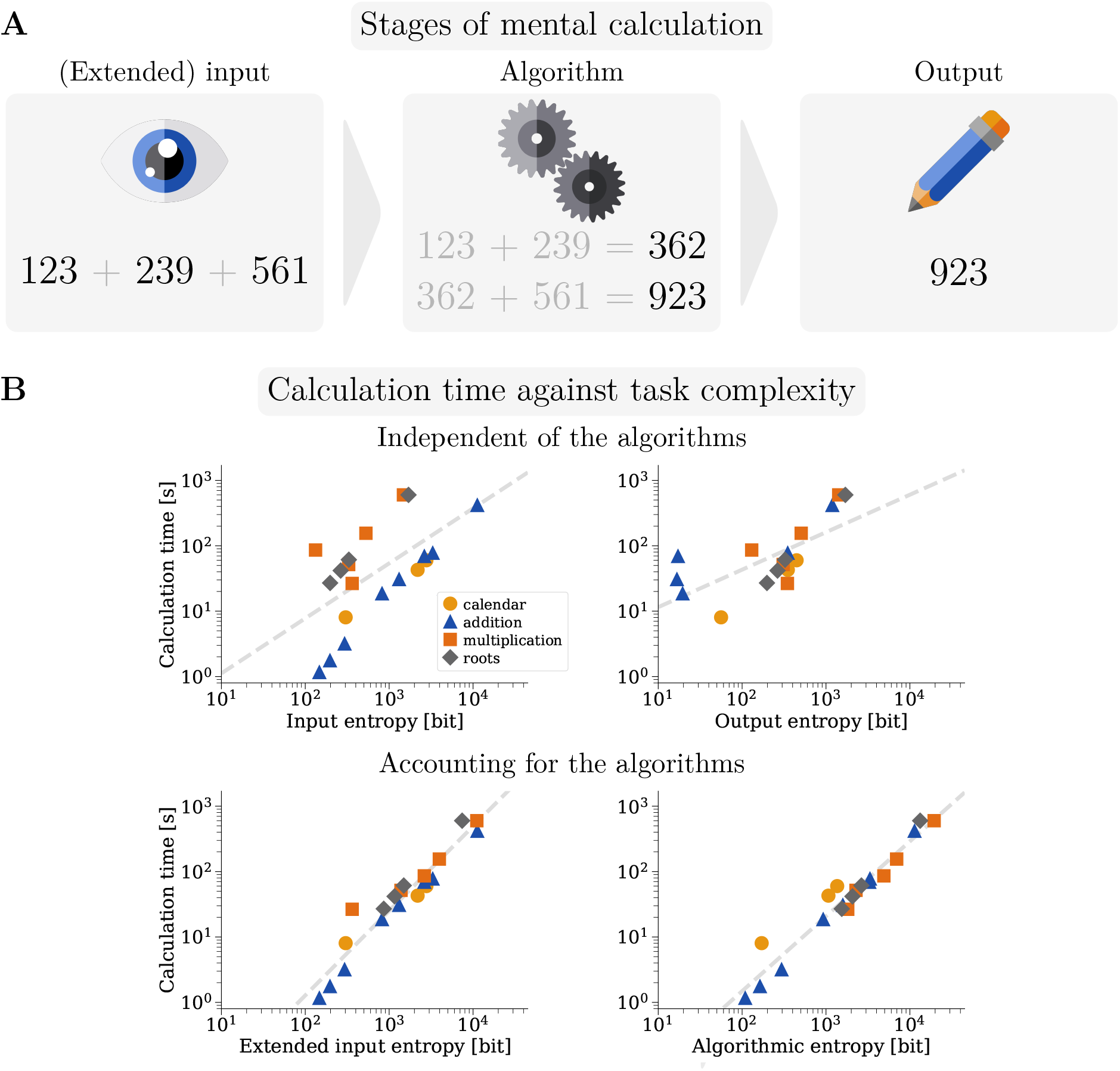
**A** Schematic of the process of mental calculation. A calculation starts with the perception of the numbers, is followed by the application of a mental algorithm, and ends with the motor output of the result. We computed entropy estimates for each of the three steps. The extended input entropy, as opposed to the raw input entropy, accounts for how many times the mental algorithm takes in the given digits or already outputted interim results. In *multiplication* and *square root* disciplines, the digits are reread several times. **B** Minimum time *T* required for mental calculation, as achieved by top-level competitors, displayed against the component entropies of the task. The top panels show the calculation time against the algorithm-independent entropies. Top left, for the input entropy, 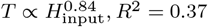 least-squares fit of logarithmic quantities, *p* = 0.0044 under t-test *H*_0_ : no correlation. Top right, for the output 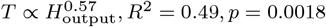 Output entropies were only computed for those disciplines in which the calculation time includes specifying the result; in the three shortest addition disciplines, the results are given after timing ended. The panels on the bottom show the calculation time as a function of entropies derived by taking the mental algorithms into account. Bottom left, as a function of extended input entropy, which counts the number of times digits, either part of the task or an already typed partial result, are read, 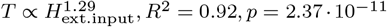 Bottom right panel, as a function of algorithmic entropy, which counts the to-be-made decisions/interim results of competitive strategies for the different tasks 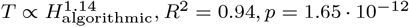 The difference between the exponents 1.29 and 1.14 is not significant (t-test for *H*_0_ : equal slopes, *p* = 0.1927).

We first computed the entropies of the input and the output across different tasks, without regard to the nature of the task or the strategy used. As a rule, calculations are performed on uniformly distributed numbers, which makes the calculation of the input entropy straightforward (see Methods). The output, though, is no longer uniformly distributed, and the entropy is generally reduced. Given that entropy, both for the input and the output, correlates with task difficulty, entropy should affect the time required to complete the tasks; indeed, there is a strong, significant correlation between calculation time and both input and output entropy (Fig. 1 **B**, top row).

Noticeably, the time needed to complete *addition* versus *square roots* calculations both scale with the complexity of the task, but distinctly. To understand why, we decided to examine the sequence of decisions successful mental calculators make. Several algorithms, specifically for the *square root* tasks and all but one of the *multiplication* tasks, reduce the memory load by reading the initial digits or interim typed or written results several times. For these tasks, we defined an extended input entropy that weights the digit entropy by the number of times the digits are read. Additionally, we estimated the algorithmic entropy of the interim results (Fig. 1 **B**, bottom row) to test whether Hick’s law applies, which states that the reaction time should scale with the entropy of possible decisions at each step^15^. The best-fit exponents for the reaction time as a function of entropy turn out to be close to unity, as predicted by Hick’s law. These exponents only skew to slightly larger values than one, with time scaling as 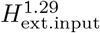 for the extended input entropy and as 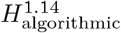 for the algorithmic entropy (t-test for the slope of the least squares fit, *H*_0_ : slope = 1, *H*_1_ : slope ≠ 1, *p* = 0.0042 resp. 0.0445).

In general, the time competitors take increases with the complexity ot the mental calculation task, whether measured in terms of input, extended input, algorithmic, or output entropy (Fig. 1 **B**). Notably, task calculation times scatter more when plotted against input or output entropy compared to when they are plotted against either extended input or algorithmic entropy. For the last two entropies, data across task categories align, forming a single, universal power-law for the calculation time as a function of the entropy. This is an indication that the cumulative entropy of the algorithmic steps is a powerful determinant of the time a mental calculation takes.

## Information rates of cognitive processing

By dividing the respective entropy by the time needed to complete a task, we can compute the processing rates for the different aspects of mental calculation (Fig. 2 **A**). The resulting ratios are conservative measures, as they assume all processes run in parallel, instead of sequentially. In general, the longer the duration of the calculations, the lower the bit rates. The extended input rate peaks at around 125 bit/s for the shortest *addition* discipline, greatly exceeding previously reported values for human information processing^1^. Across the different tasks, the algorithmic processing speed exhibits a shallower decay than the extended input rates. It starts out at almost 95 bit/s for short duration disciplines and remains above 20 bit/s even for 10-min disciplines. The smallest contribution is made by the entropy rate for the output, which remains below 15 bit/s across all tasks. The output rates exhibit almost no correlation with calculation time. To capture the total cognitive demand, we add all rates together and find that human beings hit a peak rate of around 215 bit/s over short durations, and are capable of processing 30 bit/s even for the longest disciplines (Fig. 2 **A**, bottom). These rates represent time averages instead of instantaneous values – the processing of extended input, algorithm and output might vary over the duration of each task.

**Figure 2.**
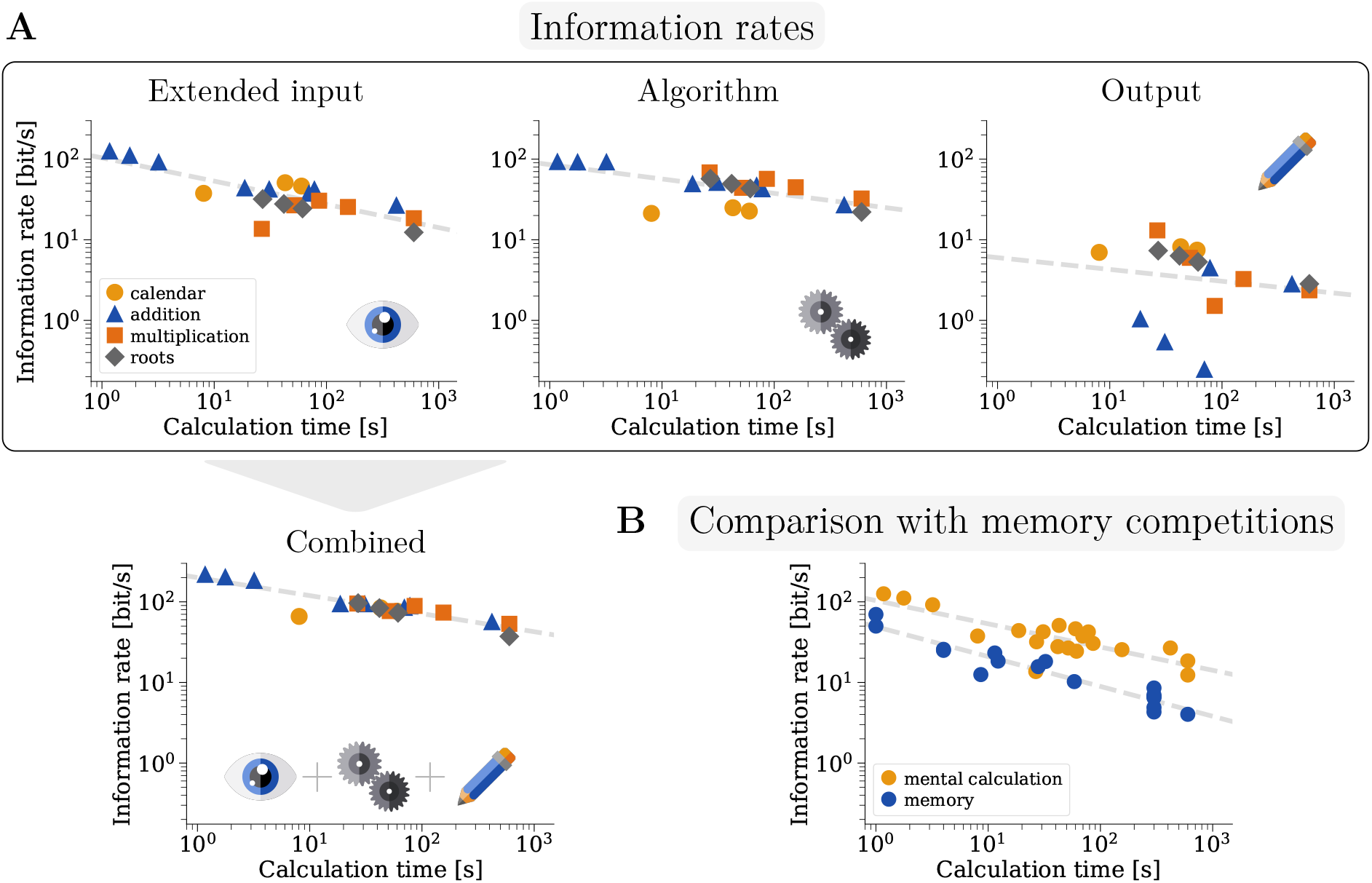
**A** Information processing rates *R* for mental calculation as a function of the calculation time *T*. Top left, extended input rates peak at over 100 bit/s for the shortest disciplines and generally stay above 10 bit/s (least-squares fit of logarithmic quantities, *R*_ext.input_(*T*) = 103.52*·* (*T/T*_0_)^*−*0.29^ bit/s, *T*_0_ = 1s, *R*^2^ = 0.66, *p* = 1.54*·*10^*−*5^ under t-test *H*_0_ : no correlation). Top middle, algorithmic information rates reach over 90 bit/s for the shortest tasks and exhibit a shallower decline (*R*_algorithmic_(*T*) = 85.66 *·*(*T/T*_0_)^*−*0.18^ bit/s, *R*^2^ = 0.43, *p* = 0.0017). Top right, output information rates remain below 15 bit/s for all tasks (*R*_output_(*T*) = 6.00 *·*(*T/T*_0_)^*−*0.15^ bit/s, *R*^2^ = 0.03, *p* = 0.53). In the three fastest disciplines, competitors indicate the result after the discipline’s timing ends. The corresponding times are, therefore, not shown in the log-log plot. The combined information rate, which sums the extended input, algorithmic and output rates, is shown as a function of the task time in the bottom left-hand panel (*R*_combined_(*T*) = 199.98*·* (*T/T*_0_)^*−*0.22^ bit/s, *R*^2^ = 0.80, *p* = 8.65 · 10^*−*8^). The combined rate peaks at over 215 bit/s for the shortest calculations and is higher than 50 bit/s in almost all disciplines. **B** Comparison of information rates achieved by mental calculation champions and memorization experts as a function of task duration, with the latter data from a recent study^14^ (upper line *R*_ext.input_(*T*) = 103.52 *·* (*T/T*_0_)^*−*0.29^ bit/s, *R*^2^ = 0.66, lower line *R*_memo_(*T*) = 49.12 *·* (*T/T*_0_)^*−*0.37^ bit/s, *R*^2^ = 0.94). The comparison is made between the extended input entropy for calculations and the input entropy for memorizations, as no estimates for the required internal processing of mnemonic techniques such as memory palaces are known. Top mental calculation performances are between two to four times faster than entropy-matched memorization performances, but the difference between the slopes of the memorization and calculation trend lines is not significant (t-test for *H*_0_ : equal slopes, *p* = 0.1455).

Many mental calculations involve a reduction in entropy from input to output, often by an order of magnitude; in contrast, memory tasks require subjects to retain the input, so that the output entropy equals the input entropy^14^. We find that results from memory competitions are consistently associated with bit rates that are several times lower than for mental calculations (Fig. 2 **B**), but the slope of time versus entropy is strikingly similar. Expert memory contestants remap the input digits to a set of “mnemonic images”^16^ resulting in a higher-dimensional, yet shorter, string, that is memorized instead. The gap between mental calculation and memory tasks is nearly constant when measured across different levels of difficulty, reflecting the higher demands encoding, consolidation, and decoding associated with memory place on cognition compared to the execution of algorithms associated with mental calculation^14^.

## Discussion

Mental calculations can proceed at more than 215 bit/s, as shown here, much faster than previously reported for cognitive tasks. Prior estimates of cognitive bit rates span a range, but are all lower. Five bit/s have been estimated for choice-reaction experiments^9^, 10 bit/s for general cognition^1^, 30*−* 60 bit/s for visual processing and attention^10,11^, 39 bit/s for human communication^12^, 50 bit/s for reading^13^, and 50*−* 70 bit/s for short memorization tasks^14^. These rates are lower by a factor of at least three compared to our findings.

The peak rate of 215 bit/s consists of 125 bit/s of visual processing and 90 bit/s of algorithmic processing. The entropy of the presented digits’ distribution is fixed, and as competitors must attend to each digit shown, the estimated visual processing rate should be reliable. In computing the algorithmic entropy, we presumed that, in some steps of the mental calculation, competitors recall memorized arithmetic facts instead of computing these from scratch; this lookup strategy saves steps in the calculation. Rudimentary arithmetic, such as knowing only the sums of single-digit numbers, would instead require many more steps and a summed entropy that would have been higher; such a minimalist assumption would have artificially inflated the algorithmic bit rate we computed. Conversely, one might ask whether competitors could gain an advantage by using larger, more extensive lookup tables; if that were the case, we might have overestimated the algorithmic bit rate. A combinatorial argument rules out this possibility, however: for instance, moving from a lookup table of the sums of two four-digit numbers to a lookup table of the sums of three four-digit numbers would imply that the human mind must be able to query 10 trillion options with a 14-digit key several times a second (see Methods).

Certainly, the individual best times of record-holders in mental calculation do not reflect average behavior or average bit rates. However, we found a remarkable regularity in the data across all studied mental calculation disciplines and across hundreds of competitors, making it less likely that our results are an accident of extremal statistics; instead, they seem to be a consequence of extensive training and high concentration shared by many contestants. Specifically, the top competitor in the short flashed addition disciplines is closely followed by the next ten competitors, who achieve speeds of at least 75% of the winning performance^17–19^.

At their best, contestants execute mental calculations at a rate equivalent to making almost one hundred yes-no decisions per second, in addition to reading in the numbers and responding with an answer. When we estimated the algorithmic bit rates, we accounted for the use of memorized partial results, tallying only those steps that resulted from willful effort. Our results show that the “speed limit” of 10 bit/s for conscious processing^20^ is regularly surpassed, even before we consider the higher bandwidth of unconscious processing.

The next question is whether these high bit rates can be sustained. Earlier studies on rapid processing built upon experiments typically lasting less than a second^21^. Mental calculation illustrates that, at least with training, high processing rates in the service of complex, conscious computation can be maintained over seconds to minutes.

The highest processing speeds in the *addition* and *multiplication* disciplines are achieved with the mental abacus technique. Here, competitors imagine the movement of the beads without the presence of a physical abacus. Previous research on abacus-trained individuals has suggested that visuomotor and visuospatial activity is involved^6,22^. This is reminiscent of participants of memory competitions, who visualize to-be-memorized information along prominent points of a familiar environment, a technique known as “memory palace”^23^. Compared to mental calculation, though, bit rates during memorization are lower by a factor of two to four (Fig. 2 **B**). One primary reason that entropy-matched memorization tasks take longer is that memory competitors map combinations of digits to predefined “mnemonic images”, such as objects or famous characters^16^. For instance, just the reading step for the memorization of digits requires more than twice as much time than a complete summation, in which only a current representation needs to be updated. Indeed, competitors take about eights seconds to read an 80-digit number for memorization^14^, compared to roughly 3.5 seconds when summing 30 three-digit numbers.

Two factors likely play a role in the striking power-law between the time to task completion and the underlying entropy. On the one hand, more complex calculations or longer sequences to be memorized are more prone to error. Performing without error in 90% of the attempts on a discipline ten times longer requires one to be error-free in≈ 99% of trials on the original discipline. Competitions, though, reward error-free performances^14^; hence competitors will adopt a more safety-oriented approach the longer and more difficult the task. On the other hand, longer task durations also make it harder to maintain the initial cognitive readiness^24^, with human subjects suffering a vigilance decrement. Clarifying these factors will require additional controlled experiments and sufficient data on the frequency with which competitors make errors.

In sum, results from mental calculators shed new light on the speed of cognition^1,20^. In general, during times of quiet contemplation, for instance, speeds of 10 bit/s might suffice. But at critical moments, we have shown that the human mind can operate at 215 bit/s or even more.

## Acknowledgments

Foremost, we are thankful for the support of the mental calculation community. Insightful comments and explanations by Samuel Engel, Marc Jornet Sanz and Daniel Timms have allowed us to improve this work substantially. Additionally, we wish to thank Wenzel Grüß, Gert Mittring, Ethan Kuntz, Aaryan Shukla and Chieko Takayanagi for conversations on mental calculation.

## Methods

### Mental calculation memory lookup versus processing

#### Arithmetic facts learned by rote

Empirically, we find that top performances of human mental calculators take a time *T* that scales non-trivially with the complexity of the task and the number of calculation steps required. Roughly speaking, if, at the *i*-th step, the set of possible numbers has size *N*_*i*_, then *T~* ∑ log_2_(*N*_*i*_). More precisely, the size of the “typical set” of numbers encountered at the *i*-th stage of calculation is 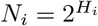, where *H*_*i*_ is the entropy.

This raises the question why competitors do not simply memorize certain arithmetic facts and then recall these. Theoretically, instead of the time *T* scaling on the order of *O*(log_2_(*N*)), memory hashes could rescale *T* to be of order *O*(1)^25^.

In fact, competitors do use arithmetic results that have been learned by rote (*personal communication*), which we factored into the algorithmic entropy computation. Take the example of adding 15 four-digit numbers flashed onto the screen, which the fastest human being can do in 1.77 seconds.

Adding 15 four-digit numbers requires 14 operations of the form

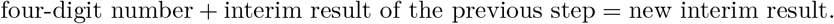

For instance, suppose that the sequence of four-digit numbers starts with 8659, 8530, … Seeing the first number 8659, there is nothing to be done, which is why there are only 14 operations to be executed. Next, 8530 has to be added to 8659, resulting in 17189, and so on. Our bit rate computations assume that each of these steps is a single, learned-by-rote operation. This requires a lookup table of size

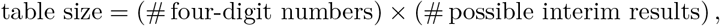

where # denotes the number of options. A four-digit number is between 1000 and 9999, which gives 9000 options. A sum of several four-digit numbers, say three, can assume values between 1000*·*3 = 3000 and 9999*·*3 = 29997. The sum of four four-digit numbers can assume values between 4000 and 39996. To cover all possible interim results, we need to consider the range from the smallest possible initial interim sum, which is 1000, to the largest possible sum of 14 four-digit numbers, which is 14*·*9999 = 139986. In total, this gives 139986*−* 1000 = 138986 options for the interim results. Hence, our current computation assumes a lookup table of size

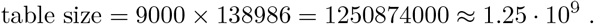

The mental calculator will be called upon to recall entries from the lookup table. If the prior likelihood of using any particular entry is uniform, then the uncertainty (or entropy) is maximal. In this case, each recall operation provides log_2_(1.25*·*10^9^) = 30.22 bit. As there are 14 operations, total algorithmic entropy would be 14*·*30.22 = 432.08 bit for this addition task. This estimate is **not** conservative, as it does not reflect the statistical regularities of the interim results: the intermediate sums are not uniformly distributed. This reduces the entropy to 207.53 bit, as explained in greater detail below. We venture no claim as to whether competitors make full or even partial use of the statistical regularities of interim results; rather, we wished to minimize the risk of overestimating the information rate. In the same spirit, for the flashed addition tasks, we excluded operations that could be carried out after the presentation sequence ends, leaving only 163.24 bit of algorithmic entropy remaining for the addition of 15 four-digit numbers (please see the section on addition).

In sum, when competitors use a lookup table for four-digit additions, with each such addition treated as a single step in the calculation, their algorithmic rate surpasses 90 bit/s, at the very least (see Table 1 for the results). This lower bound is derived by dividing 163.24 bit by the 1.77 second-long duration in which 15 four-digit numbers are presented to the fastest competitors.

**Table 1.**
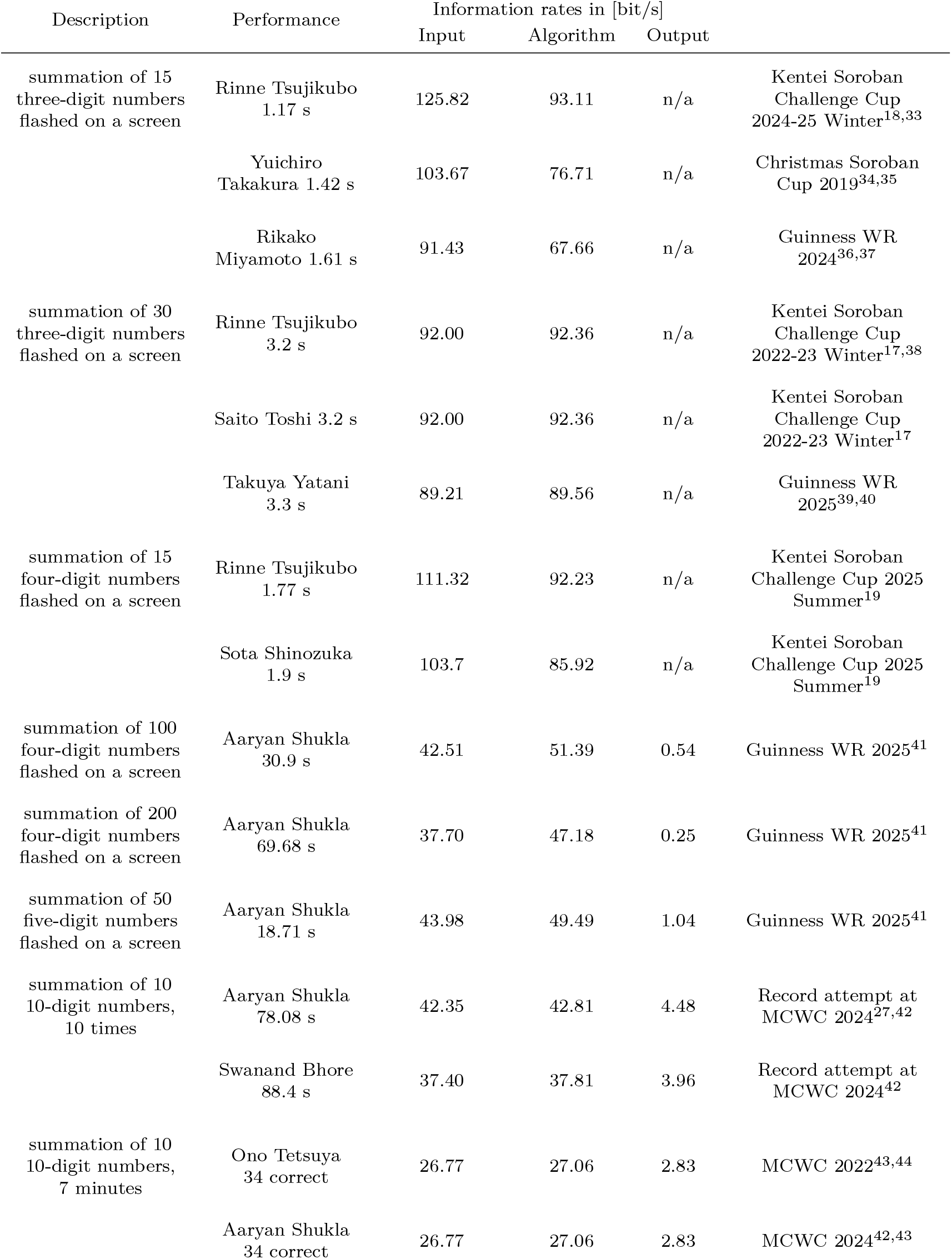
Performances of top competitors in mental *addition* disciplines. If the time to give the result is not included in the record, the output rate is listed as n/a.

**Table 2.** Performances of top competitors in mental *multiplication* disciplines.

| Description | Performance | Information rates in [bit/s] |  |  |  |  |
| --- | --- | --- | --- | --- | --- | --- |
|  |  | Input | Ext.<br>Input | Alg. | Output |  |
| multiplication of a six and a five-digit number, 10 times | Hiroaki Tsuchiya<br>26.54 s | 13.65 | 13.65 | 68.43 | 13.06 | Osaka Soroban Championship 2024<sup>45</sup> |
| multiplication of two five-digit numbers, 10 times | Aaryan Shukla<br>51.69 s | 6.37 | 26.76 | 44.10 | 5.97 | Guinness WR 2025<sup>41,46</sup> |
|  | Marc Jornet Sanz<br>59.4 s | 5.54 | 23.28 | 38.37 | 5.20 | Record attempt 2019<sup>46</sup> |
| multiplication of two eight-digit numbers, 10 times | Aaryan Shukla<br>155.41 s | 3.40 | 25.51 | 44.76 | 3.26 | Guinness WR 2025<sup>41,46</sup> |
|  | Marc Jornet Sanz<br>162.3 s | 3.26 | 24.43 | 42.86 | 3.11 | Record attempt 2019<sup>46</sup> |
| multiplication of two eight-digit numbers, 10 minutes | Aaryan Shukla<br>28 correct | 2.47 | 18.50 | 32.46 | 2.36 | MCWC 2024<sup>42,43</sup> |
|  | Tomohiro Iseda<br>25 correct | 2.20 | 16.52 | 28.99 | 2.11 | MCWC 2018<sup>43,47</sup> |
| multiplication of two 20-digit numbers | Aaryan Shukla<br>86 s | 1.54 | 30.52 | 56.93 | 1.53 | Unofficial record attempt 2024<sup>48</sup> |
|  | Aaryan Shukla<br>105 s | 1.26 | 25.00 | 46.62 | 1.24 | Record attempt at MCWC 2022<sup>49</sup> |
|  | Grant Thakkar<br>168 s | 0.79 | 15.63 | 29.14 | 0.77 | Record attempt at Memoriad 2014<sup>46</sup> |

**Table 3.** Performances of top competitors in *calendar* calculations.

| Description | Performance | Information rates in [bit/s] |  |  |  |
| --- | --- | --- | --- | --- | --- |
|  |  | Input | Algorithm | Output |  |
| 125 random dates from 1600 – 2100 | Aaryan Shukla<br>42.84 s | 51.00 | 24.95 | 8.19 | Unofficial record attempt 2024<sup>50</sup> |
|  | Jay Baldiya Jain<br>56.82 s | 38.46 | 18.81 | 6.18 | Memoriad 2016<sup>46,51</sup> |
| from 1600 – 2100, one minute, as many dates as possible | Yusnier Viera<br>159 correct | 46.32 | 22.66 | 7.44 | Record attempt 2024<sup>46</sup> |

**Table 4.** Performances of top competitors in mentally extracting square roots of six-, eightand 10-digit numbers.

| Description | Performance | Information rates in [bit/s] |  |  |  |  |
| --- | --- | --- | --- | --- | --- | --- |
|  |  | Input | Ext.<br>Input | Alg. | Output |  |
| square roots of six-digit numbers to five decimal places, 10 times | Aaryan Shukla<br>26.99 s | 7.33 | 31.94 | 57.1 | 7.33 | Unofficial record attempt 2024<sup>52</sup> |
|  | Shashank Jain<br>27.3 s | 7.25 | 31.58 | 56.45 | 7.25 | Record attempt at Memoriad 2024<sup>46,53</sup> |
| square roots of six-digit numbers to five decimal places, 10 minutes | Aaryan Shukla<br>86 correct | 2.84 | 12.36 | 22.09 | 2.84 | MCWC 2024<sup>42,43</sup> |
|  | Kaloyan Geshev<br>69 correct | 2.27 | 9.92 | 17.72 | 2.27 | MCWC 2024<sup>42,43</sup> |
| square roots of eight-digit numbers to five decimal places, 10 times | Aaryan Shukla<br>41.77 s | 6.33 | 27.80 | 49.51 | 6.33 | Unofficial record attempt 2024<sup>54</sup> |
| square roots of 10-digit numbers to five decimal places, 10 times | Aaryan Shukla<br>61.18 s | 5.40 | 24.41 | 43.22 | 5.31 | Unofficial record attempt 2023<sup>55</sup> |

Similar considerations will apply to other mental calculation dsiciplines. Next, we address the question of whether the lookup tables that competitors have learned by rote are reasonably sized.

### Combining calculation steps in even larger lookup tables is implausible

Were competitors to summarize several addition steps into a single step, the required lookup table would increase exponentially in size. To demonstrate the combinatorial explosion, we return to the example of adding 15 four-digit numbers. We compute the size of the lookup table that will permit one to add any pair of four-digit numbers to any possible interim result.

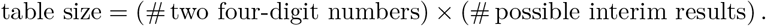

There are 9000^2^ options for two four-digit numbers. Possible interim results are the sums of one, three, five, seven, nine, eleven or thirteen four-digit numbers. The one four-digit numbers is at least 1000 and the sum of 13 four-digit numbers is at most 13 *·*9999 = 127987. Therefore, one would need to cover

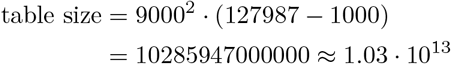

combinations, each accessible with an up to 14-digit key (two four-digit numbers and a six-digit interim result). Given that the best calculators see the 15 four-digit numbers for less than 2 s, they would need to distinguish the 10 trillion options with the 14-digit key repeatedly within that time.

We conclude that mental calculations pass through a set of interim results that only require reasonably sized memorized lookup tables. (Moreover, in abacus competitions, the numbers are flashed in succession, so contestants cannot even add columns of multiple digits all at once.)

### Entropy calculations

We first introduce input, extended input, algorithmic and output entropy in general, before computing these quantities for each discipline. We measure entropy by the Shannon formula

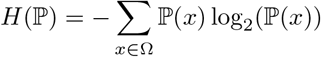

for a distribution ℙ on a discrete probability space Ω. An outline of the entropy of mental calculations has previously been given by Timms^26^.

#### Input

We consider the entropy of all numbers on which the calculations are performed. In most tasks, these are uniformly drawn *k*-digit numbers with entropy

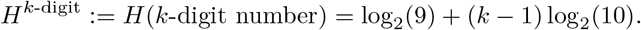

as the first digit is non-zero. For example, the input entropy of any operation on two uniform single-digit numbers, e.g., their sum, is 2*H*^1-digit^ = 2 · log_2_(9).

#### Extended input

To reduce the load on working memory, various mental calculation algorithms involve reading either the presented digits or already written or typed interim results several times. The extended input entropy accounts for these digits.

Only when the task is parsed exactly once do the input and extended input coincide. This is the case for *addition, calendar dates* and one of the *multiplication* disciplines.

Otherwise, the entropy for the extended input is larger than for the input. One *multiplication* algorithm involves reading each input digit several times. In the *square roots* disciplines, competitors use the already calculated digits for subsequent iterations of the algorithm.

#### Algorithm

To determine the entropy of the algorithm, we sum the entropies of the “decisions”, which are interim outcomes along the steps of the algorithm. For instance, when adding two uniform, nonzero, single-digit numbers, different input combinations result in the same sum, e.g. 3 + 2 = 5 = 1 + 4, leading to 17 combinations in the outcome space that are not equally distributed. The entropy of this distribution is 3.88 bit. During the summation of three uniform single-digit numbers, we consider the two distributions that arise: first, the sum of two uniform single-digit numbers, and then the sum of the resulting number and another uniform single-digit number. We add the entropies of both distributions to estimate the algorithmic entropy.

The employed strategies are not minimal in terms of algorithmic entropy. Even though the number of digits of expert calculators store in short-term memory are typically larger than average^5^, working memory is a major limitation in mental calculation, as it restricts the nature of the algorithms easily executed by human competitors. An example are multiplications, where two strategies are commonly in use. The first requires less entropy processing, but the working memory load grows quickly with the number of digits, whereas the second strategy requires more entropy processing, but places little demand on working memory. Consequently, the first strategy yields superior results for manageable multiplications, whereas the second is applied to large multiplications such as the product of two 20-digit numbers.

#### Output

The numbers that the calculator must type or write down define the output entropy. We do not account for the coordination of multiple muscles and prompt correction of errant hand trajectories^20^, so this output entropy is a lower bound on the entropy of the reporting process.

In many disciplines, competitors type or write the results continuously throughout the task, not only at the end. The reason is two-fold: either many instances of the same elementary exercise (e.g. *calendar* calculations) are part of the task, or the algorithm successively produces new digits of the result (e.g. *multiplication*). Shorter-duration disciplines work differently, however. In the flashed *addition* disciplines, for example the summation of 100 four-digit numbers, competitors present the result only at the end. In the three shortest *addition* disciplines, the completion time does not include reporting the result; in these cases, we discount the output entropy in the sum of entropies, artificially setting the output entropy to zero, which ensures that the summed entropy rates still represents a lower bound on the true information rate.

Human competitors might not be able to make full use of the knowledge of the distribution of the calculation results. The output entropy can be considered a lower bound on the “perceived” entropy.

### Addition

Competitors add uniformly distributed numbers, which are either flashed at a certain rhythm or shown all at once.

#### Input

If there are *n* uniform, *k*-digit numbers to be added, the input entropy is given by *nH*^*k*-digit^. In the flashed *addition* disciplines, numbers are replaced on-screen 2.5 to 13 times per second.

#### Algorithm

Experts employ the mental abacus technique: after years of training the calculations, competitors are able to imagine the movement of the beads without the presence of a physical abacus^6^. As mental calculators pass through a set of interim sums, we generalize the initial addition examples to sums of multi-digit numbers. To this end, let us introduce the notation

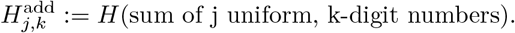

For instance, for the addition of two uniform four-digit numbers, the resulting distribution on 2000, 2001, … up to 19998 is used to compute the associated entropy. We obtained the following approximate values for the addition of two numbers

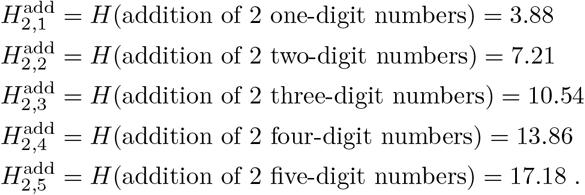

An addition of two numbers with several digits could alternatively occur by adding single-digit numbers successively. Interestingly, the above values indicate that the entropy is larger using single-digits than adding several digits at once, as there are more decisions to be made using single-digits:

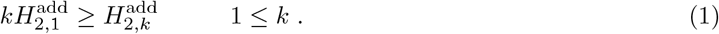

This might explain an observation from the “summation of 10 10-digit numbers” discipline, where all numbers are shown at once: top competitors choose to sum columns containing several digits, typically two four-digit and a final two-digit column^27^. A proof of the inequality (1) is given at the end of the entropy calculation section.

For a summation of several numbers, we add the entropies of all the intermediate distributions

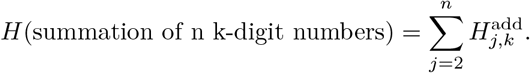

For the three shortest disciplines, the summation of 15 three-digit, 15 four-digit and 30 three-digit numbers, this formula results in

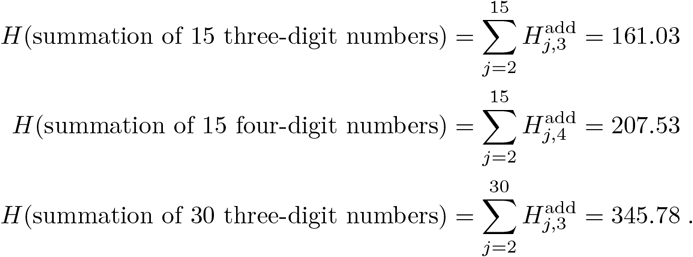

In these disciplines, the timed duration only includes the flashed presentation of the digits, but not the time used to report the result. The differences between the respective world records of 1.17, 1.78 and 3.2 s by Rinne Tsujikubo allow us to estimate to what extent the calculations are carried out after the timing ends. For the respective disciplines, let *e*_*b,i*_ be the entropy processed before and *e*_*a,i*_ be entropy processed after the flashed presentation ends. Let us assume that the rate at which calculations are carried out is similar, whether the discipline lasts one or three seconds, i.e.,

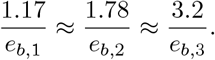

If we posit that the amount of entropy processed after the timing ends is similar across the three disciplines *e*_*a*,1_ = *e*_*a*,2_ = *e*_*a*,3_ =: *e*_*a*_, we obtain a second system of equations

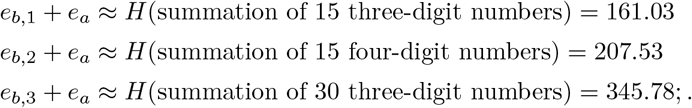

Solving the full, overdetermined system with least squares yields

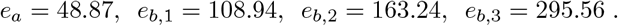

The entropy *e*_*a*_ corresponds to about four addition operations. We use *e*_*b,i*_ to calculate the algorithmic rates for the three disciplines.

The summed algorithmic entropies are

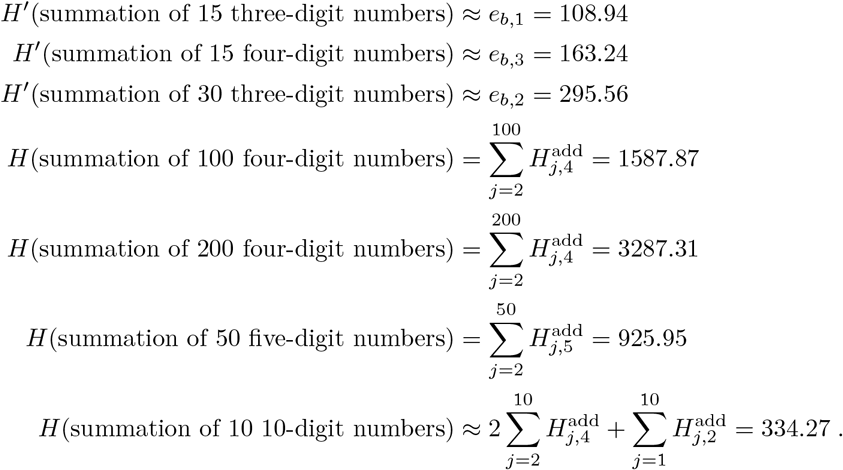

#### Output

We computed

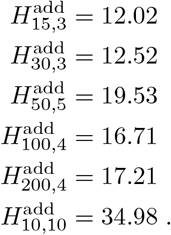

Indeed, these values are consistent with with the prediction from the central limit theorem

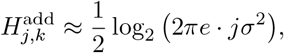

where *σ*^2^ is the variance for a uniform distribution between 10^*k−*1^ and 10^*k*^ *−* 1:

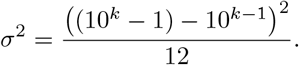

### Multiplication

Competitors multiply uniformly distributed, integer numbers with *k*_1_ and *k*_2_ digits, where *k*_1_, *k*_2_ are at least 5.

#### Input

The number of solved tasks was multiplied by 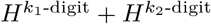 to determine the entropy.

#### Algorithm

We discuss two methods. The first “traditional method” requires the ability to visualize several digits simultaneously, achievable by a mental abacus, whereas the second “cross-multiplication method” places little demand on visualization and working memory^28^. The traditional method is superior in terms of entropy. However, as the cross-multiplication places a limited burden on working memory, it generalizes to almost arbitrarily large numbers, for instance the multiplication of 20-digit numbers, where the traditional method becomes infeasible.

##### 1. Traditional method

This approach is used by top competitors at Japanese soroban competitions^28^. The mental algorithm is equivalent to the pen-and-paper method one might have learned in school. For example, to multiply 323 *·* 446, one performs three multiplications of 4, 4, 6 with 323 and adds the results:

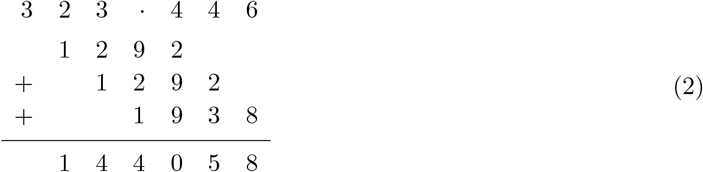

To understand the entropy of a six-digit by five-digit multiplication, note that there will be six multiplications of a single-digit by a five-digit number and five five-digit additions. As visible from (2), if consecutive numbers in the right-hand factor are equal, one of the singleby six-digit multiplication repeats and we do not count its entropy. The probability for this event is 0.1 and, as the first digit cannot repeat, we assume that there are 6 *−* 5 *·* 0.1 = 5.5 multiplications on average. We computed

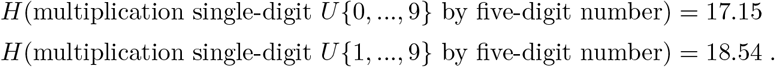

In total, this led us to the formula

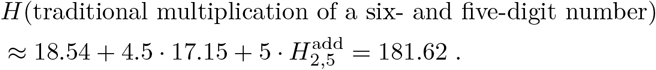

##### 2. Cross-multiplication method

This strategy is more common at the MCWC^29^. The method builds upon the idea to collect all terms that contribute to a certain power of 10. For example, the product 323*·* 446 is calculated by collecting all terms associated to 1, 10, 100, …, which can be visualized through an eponymous pattern of crosses:

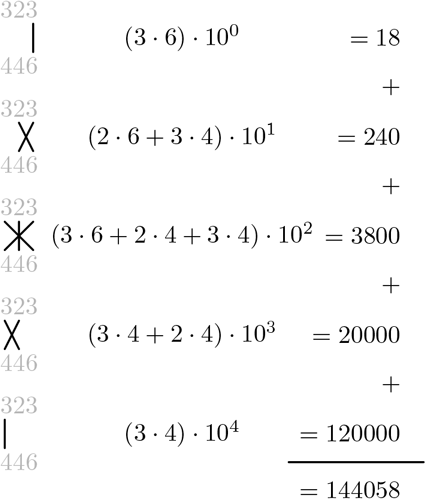

By carrying out the operation from right to left, one automatically keeps track of the current power of 10. If the number of digits *k*_1_ and *k*_2_ of the two factors differ, suppose *k*_1_ *< k*_2_, zero can be appended *k*_2_ *−k*_1_ times to the beginning of the smaller number. For this reason, and as all mental calculation results stemming from this method involve numbers of equal length, let *k*_1_ = *k*_2_ = *k*.

We assumed that each of the digits of the factors is uniformly distributed on *{*0, …, 9*}* except the leading digit, which is uniformly distributed on *{*1, …, 9*}*. Generalizing the above algorithm to the multiplication of *k*-digit numbers, there will be 2*k−* 1 steps, involving the addition of 1, 2, 3, …, *k−* 1, *k, k −*1, …, 3, 2, 1 times the product of two uniform single-digit numbers. We carried the exact intermediate distribution along all of these steps, accumulating the entropy for each of the additions and multiplications to arrive at

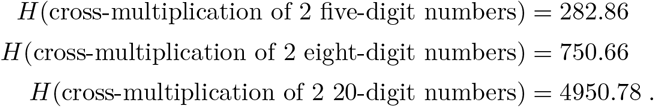

As top competitors have memorized the results of multiplying any two two-digit numbers, which occur at the begin and end of the algorithm, we subtracted

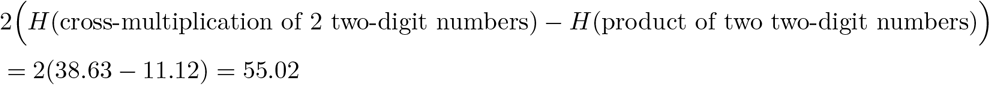

from each of the above values.

#### Extended input

##### 1. Traditional method

Abacus experts can perform this computation after hearing the numbers, suggesting that no rereading is necessary.

##### 2. Cross-multiplication method

The leading non-zero digits are each read *k* times. For the remaining non-zero digits, 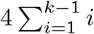 reads take place. Memorizing two-digit products avoids one reading each of the digits, leading to eight fewer reads. This translates to the expression

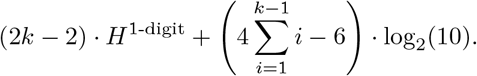

#### Output

Multiplication of integers commutes, with *n ·m* = *m ·n*, so each result that is not a perfect square appears at least twice in an enumeration of all products. Moreover, depending on the prime factorizations of *m* and *n*, a number of products appear more times. If we ignore the multiplicities, we have a simple upper bound

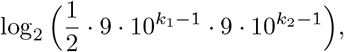

which arises from the fact that there are 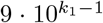 and 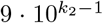 options for the two factors and the number of distinct products is lower by a factor of at least two.

Enumeration of all products quickly becomes intractable, so we resorted to Monte-Carlo sampling of the products to estimate the average inverse multiplicity, while respecting the bounds on the range of integers. We sampled 10^6^ products *n· m* and computed the number of combinations of *n* and *m* that yield the same product, but also lie within the desired range, i.e., have the required *k*_1_ and *k*_2_ decimal digits. Denoting this number as *µ*(*n ·m*), we computed the average inverse ⟨*µ*^*−*1^⟩ and extrapolated from 10^6^ to the number of possible products. This then yielded

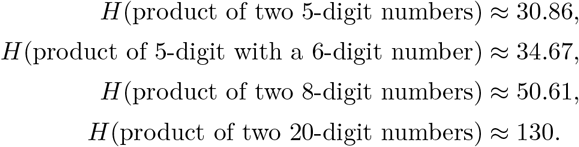

Compared to the input entropy, the multiplicities above reduced the number of distinct products by a factor of roughly four to six.

### Calendar calculations

Participants are given calendar dates and years and must determine the corresponding days of the week.

#### Input

If the discipline features years between [*x*_1_, *x*_2_), the entropy of a single date is

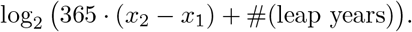

#### Algorithm

The Gregorian calendar repeats every 400 years, so that August 13, 2025, and August 13, 2425, fall on a Wednesday, for instance. The most common competition format spans the years 1600 2100 and does not include years prior to the 17th century, as the Gregorian calendar came into effect in 1582. In one 400-year cycle, 97 are leap years,

Calendar calculations correspond to an addition modulo seven. Dates are divided into century, (e.g. 19-th), year (e.g. 84), month and day, each represented modulo seven, with a leap-year correction to the day in the months of January and February..

Instead of adding four separate summands for century, year, month and day, top competitors have memorized the 400 joint century/year combinations and 365 joint day/month combinations leading to a single modular addition^30^. The century/year and day/month contributions are essentially uniformly distributed, each amounting to a decision with entropy log_2_(7). The addition modulo 7 carries another log_2_(7) bit. The two months January and February constitute

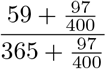

of an average year, which is why we add

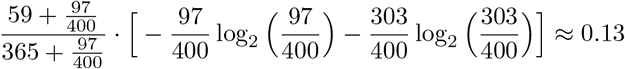

for a leap-year rule. We have numerically checked that the first two decimal places of the value are the same for the year ranges used in the two calendar date disciplines.

Whether competitors deduce the day of the week from dates in the current century or from the years 1600-2100, the same algorithm applies. We assign an algorithmic entropy per unit computation of

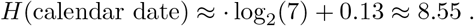

#### Output

The output is one of the seven options Monday, Tuesday, Wednesday, Thursday, Friday, Saturday, Sunday, so log_2_(7) per date.

### Square roots

Competitors extract roots of six-digit, eight-digit and ten-digit numbers to five decimal places. Numbers with integer roots are excluded to guarantee non-trivial decimal places.

#### Input

We computed the entropy by multiplying the values *H*^6-digit^, *H*^8-digit^ and *H*^10-digit^ with the number of correctly extracted square roots.

#### Algorithm

A common choice among top competitors is the duplex method^29,31^. We use 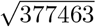 as an example to illustrate the method. Partial results are indicated by bold digits. The first digit of 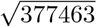 can be determined by considering only the first two digits and searching for the largest single-digit square, which is smaller, here **6** as 36 *<* 37. Subtracting 36 from the first two digits results in a new number 17463. To generate the additional decimal places, the following steps are repeated:

1. Divide the first two digits by twice the first digit, here 17*/*(2 *·*6) = **1** remainder 5. Append the next digit of 17463, i.e., 4 to the remainder, resulting in 54. If no digits are left, append zero instead.
2. Form the difference of this number and the duplex of the interim result, ignoring the interim’s first digit. The duplex of a *k*-digit number *a*_1_*a*_2_…*a*_*k*_ is defined as

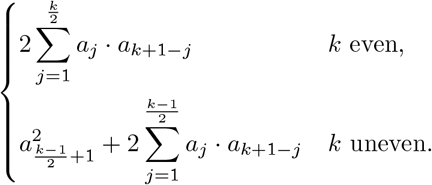

In essence, one multiplies the first and last digit, the second and second-to-last digit … and takes twice the sum of all these products. If *k* is uneven, the resulting middle digit is squared and added to the sum. In our case, this results in 54 *−* 1^2^ = 53.

If the result after step 2. is negative, revert to 1. and consider an improper remainder instead. Iterating the above steps, we come across this case at 48*/*12 = **3** and 27*/*12 = **1**:

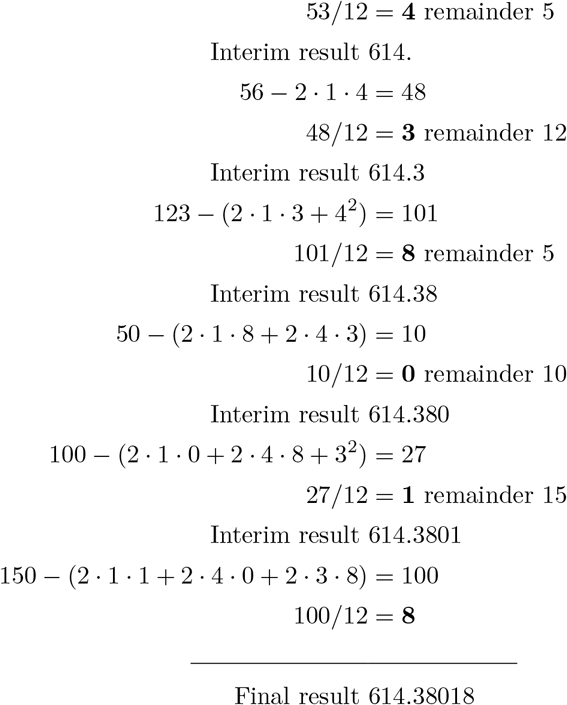

Participants can and do memorize the squares from 1 to 999 to estimate the first two digits of the result, hereby skipping the first two iterations^5^. In the example above, **614**^2^ *<* 377463 *<* 615^2^. Given the task of taking the square root of a six-digit number, the integer part of the result will lie between 316 and 999 and have entropy 9.35. Divisions then decide the next digit of the final result, a log_2_(10) decision, leaving a remainder, for which the number of possibilities amounts to two times the first digit *X* of the square root. Assuming that the remainders are uniformly distributed, and ignoring improper remainders, we computed the entropy of the remainder decision to be E(log_2_(2 *·X*)) = 3.67. To calculate the entropy of the duplexes, let

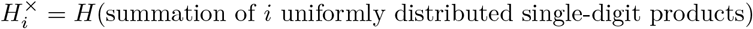

and, introducing the term “half square” for the square term of odd-digit duplexes,

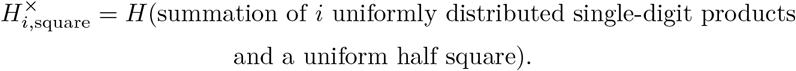

As a square of a single-digit is simple, we set

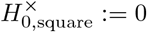

and computed the following values

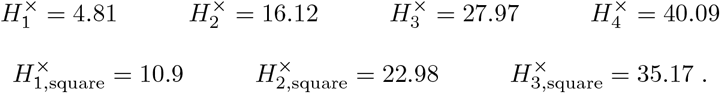

To compute *H*^*×*^, we counted each of the single-digit products and computed each of the additions similar to the approach for the *addition* disciplines. In a majority of cases, a duplex is a two-digit number, and if more than four uniform digits *a*_1_*a*_2_*a*_3_*a*_4_ are involved, even a three-digit number. Therefore, as we skip the first two iterations, we accounted for the subtractions of the four remaining duplexes by two two-digit subtractions of 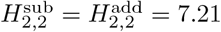 bit and two three-digit subtractions of 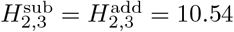 bit.

Adding all the entropies, we arrived at

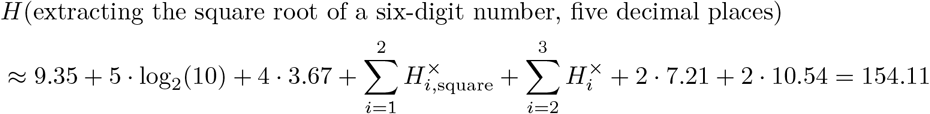

The same algorithm generalizes to the square roots of eight-digit and 10-digit numbers to five-decimal places. We require one or two additional iterations, consisting of a remainder operation with log_2_(10) + 3.67 = 6.99 bit and additional subtractions of duplexes. This allows us to write

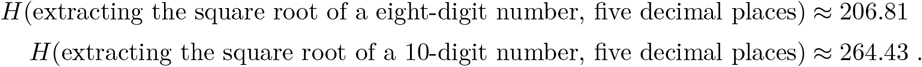

#### Extended input

The algorithm shows that, after the initial step, which generates three digits of the result, competitors reread one, then two, three, four… of the already calculated digits to generate the next digit. As the first iterations can be skipped through memorization, we added 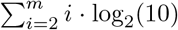 to the input entropy, where *m* = 6, 7, 8.

#### Output

The mapping *x⟼* round(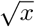, 5) is one-to-one for six- and eight-digit numbers, and the output entropy agrees with the input entropy. At most two ten-digit numbers are mapped to the same output; the output entropy was computed to be 32.51 compared with the input entropy 33.07 = *H*^10-digit^.

### Proof of the entropy inequality

This subsection is dedicated to the proof of (1). To deduce the statement, note the monotone convergence

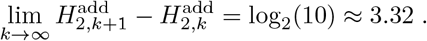

Inductively, this entails 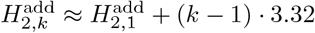 and as 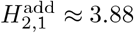, we get 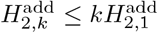.

To show the above limit, we will first derive the distribution of the sum of two uniform *k*-digit numbers and then a formula for the associated entropy 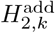.

For instance, the sum of two single-digit numbers is between 2 and 18. There is one combination to obtain 2 = 1 + 1, there are two combinations to obtain 3 = 2 + 1 or 3 = 1 + 2, three combinations for 4 and so on. For a sum of 10, there are nine combinations and from there on the number of combinations decreases. As any of the combinations is equally likely with probability 1*/*9^2^ = 1*/*81, the complete distribution is given by

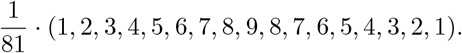

Proceeding in the same way for *k*-digit numbers, where the probability for any one combination is (9 *·* 10^*k−*1^)^*−*2^ and the number of combinationsd increases to 9 *·* 10^*k−*1^ before decreasing, the distribution of the sum is

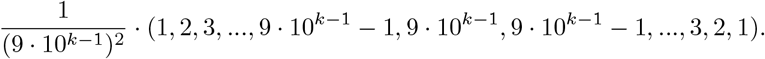

The entropy of this distribution is given by

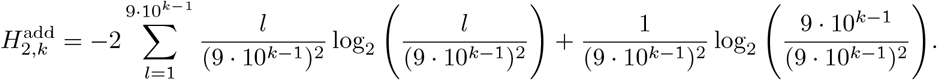

Setting *C*_*k*_ := 9 *·* 10^*k−*1^ and rewriting the second term, this expression becomes

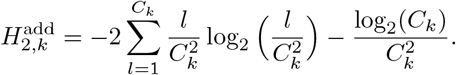

The sum can be approximated by a Riemann-integral

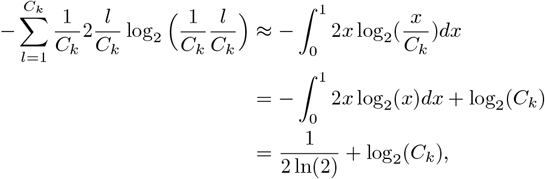

which yields

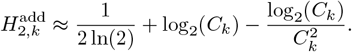

As *C*_*k*+1_ = 10*C*_*k*_ and 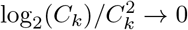 as *k ⟼ ∞*, the convergence follows from log_2_(*C*_*k*+1_) = log_2_(10) + log_2_(*C*_*k*_).

## Author contributions

BW conceived the study and performed the data analysis. All authors contributed to the conceptual analysis, writing, and editing of the manuscript.

## Competing interests

The authors declare no competing interests.

## Data availability

All considered data is based on public references and listed in the extended data section.

## Extended data

We used public sources to compile the lists of mental calculation performances in Table 1-4. We focused on mental calculators who successfully participated in major mental calculation competitions, primarily the Mental Calculation World Cup (MCWC) and the Japanese soroban/abacus competitions. If several references are listed for the same discipline, the best result is shown in Fig. 1 and Fig. 2.

Not all competition formats have been included in this analysis. For example, several world records involving finding the exact roots of larger numbers such as the 13-th root from a 100-digit number do not lend themselves to our entropy-based analysis. These records rely on intricate algorithms and occasionally fortunate combinations that make entropy estimation infeasible^3,32^.

## Notes

### Competing Interest Statement

The authors have declared no competing interest.

### Summary of Updates

Improved representation, figures, text and calculations. Added a section on lookup tables.

